# Evidence for an one-step mechanism of endosymbiont-induced thelytoky in the parasitoid wasp, *Muscidifurax uniraptor*

**DOI:** 10.1101/2022.10.27.514028

**Authors:** Yidong Wang, Eveline C. Verhulst

## Abstract

*Wolbachia* manipulates host reproduction in various haplodiploid insect species, in which fertilized eggs normally develop into diploid females while unfertilized eggs develop into haploid males. Females infected with a thelytoky-inducing *Wolbachia* produce diploid daughters from unfertilized eggs (thelytoky), but in some infected species diploid males spontaneously occur in low numbers. This suggests that diploidization and feminization are induced separately. In the *Wolbachia-*infected thelytokous parasitoid wasp, *Muscidifurax uniraptor*, occasional males have been found but with unknown ploidy. Therefore, we studied the mechanism of *Wolbachia*-induced thelytoky in *M. uniraptor* in the context of sex determination. We started by feeding different concentrations of tetracycline (antibiotic) to *M. uniraptor* females to gradually reduce the *Wolbachia* titre. We found that a decreased *Wolbachia* titre leads to an increased proportion of haploid male offspring, but we found no diploid males. Next, we studied the effect of *Wolbachia* infection on the expression and splicing of the sex determination genes *transformer* (*Mutra*) and *transformer-2* (*Mutra2*) in female ovaries and conclude that *Wolbachia* does neither affect the expression nor the splicing of *Mutra* and *Mutra2*. We then measured *Mutra* and *Mutra2* expression levels in developing zygotes at different time points and found a two-fold *Mutra* expression increase in *Wol*+ compared to *Wol*-. Finally, we used the sexually reproducing sister species, *Muscidifurax raptorellus* and artificially created triploid females to determine whether diploidization is sufficient for feminization. These triploid females, when virgin, produced haploid sons and diploid daughters, showing that in *Muscidifurax* feminization solely depends on ploidy level. This strongly suggests that *Wolbachia* only needs to induce diploidization and that bi-allelic *Mutra* expression is sufficient for female development.

## Introduction

All insects belonging to the order Hymenoptera have a haplodiploid mode of reproduction, whereby fertilized eggs develop into diploid females and unfertilized eggs develop into haploid males (Cook, 1993). However, in a small subset of haplodiploid species, unfertilized eggs develop into females in a process termed thelytoky. This type of parthenogenesis can be induced by nuclear genes (Beukeboom and Pijnacker, 2000; Lattorff et al., 2005; Rössler and DeBach, 1973; Schilder et al., 1999; Vavre et al., 2004; Verma and Ruttner, 1983) or by intracellular endosymbionts, such as *Wolbachia*, *Rickettsia* and *Cardinium* (Chigira and Miura, 2005; Furihata et al., 2015; Hagimori et al., 2006; Stouthamer et al., 1990; Zchori-Fein et al., 2004). These endosymbionts are vertically transmitted to the offspring only through the egg cytoplasm, as sperm cells do not contain enough cytoplasm (Ma and Schwander, 2017). By increasing the ratio of infected females in the population through endosymbiont-inducted thelytoky, they obtain a selective advantage. It is assumed that endosymbiont-induced thelytoky is only present in host species with haplodiploid sex determination as egg development can commence without fertilization (Leach et al., 2009).

Realizing diploidization of haploid eggs seems a prerequisite for endosymbionts to induce thelytoky in a haplodiploid reproduction system, as feminization would follow automatically from the host’s haplodiploid mode of reproduction (Kageyama et al., 2012; Werren et al., 2008). However, in some endosymbiont-induced thelytokous species it was found that lowering the endosymbiont titre by antibiotic curing or heat treatment resulted in diploid male progeny, and only complete endosymbiont removal leads to haploid male progeny (Ma et al., 2015; Tulgetske, 2010). This suggests that a low endosymbiont titre is sufficient for diploidization and a higher endosymbiont titre is required for feminization, essentially decoupling diploidization and feminization from each other in a two-step mechanism (Ma et al. 2015).

The mechanisms by which endosymbionts would induce feminization in its host are unknown, and it is further complicated by the extraordinary diversity of hymenopteran sex determination mechanisms (Heimpel and de Boer, 2008). In all Hymenoptera studied to date, the splicing of terminal sexual differentiator, *Doublesex (Dsx)*, is controlled by the upstream splicing regulators Transformer (Tra) or its ortholog Feminizer (Fem), and Transformer-2 (Tra2) (Gempe et al., 2009; Geuverink et al., 2017; Hasselmann et al., 2008; Inoue et al., 1992; Nissen et al., 2012; Verhulst et al., 2010a). To guarantee female offspring development in Hymenoptera, it is essential to maintain the female-specific splicing of *tra* (or *fem*). In the parasitoid wasp *Nasonia vitripennis*, *tra* and *tra2* are maternally provided to the eggs to initiate female-specific splicing of early zygotic *tra* messenger RNA (mRNA) (Geuverink et al., 2017; Verhulst et al., 2010a). The continuous splicing of female-specific *tra* (or *fem*) mRNA is regulated by its own protein, leading to female development (Geuverink et al., 2017; Verhulst et al., 2010a). Recently, it has been shown in *N. vitripennis* that an active copy of *wasp overruler of masculinization* (*wom*) from the paternal genome is also required for zygotic *tra* expression in order to start its autoregulation and female development (Beukeboom and van de Zande, 2010; Verhulst et al., 2010a; Verhulst et al., 2013; Zou et al., 2020). An inactive, imprinted *wom* copy from only the maternal allele does not lead to sufficient zygotic *tra* expression, resulting in male development (Zou et al., 2020). In the honey bee, *Apis mellifera*, heterozygosity at the *complementary sex determiner* (*csd*) locus and *tra2* expression result in initial female-specific *fem* splicing (Beye et al., 2003; Hasselmann et al., 2008; Nissen et al., 2012). Then, a positive feedback loop of female-specific *fem* splicing is mediated by its own Fem protein (Gempe et al., 2009). Hemi-or homozygosity at the *csd* locus results in male development (Beye et al., 2003). In *Asobara tabida*, although the exact initial signal leading to female-specific *tra* autoregulation is still unknown, maternal provision of *tra* and *tra2* are presumed to be important for female development (Geuverink et al., 2018). The ubiquity of *tra2* and female-specific *tra*/*fem* and their crucial function in the female sex determination cascades, offers a potential for endosymbionts to manipulate their early expression in order to induce feminization.

In the *Wolbachia*-infected thelytokous wasp, *Muscidifurax uniraptor*, spontaneous males are occasionally produced by *M. uniraptor* females in laboratory rearing. It was shown previously that curing females from *Wolbachia* infection using antibiotics results in male offspring production, and an increasing dose of antibiotics results in a higher frequency of male occurrence (Zchori-Fein et al., 2000). However, diploid males have not been recorded in literature, and ploidy testing of the spontaneous males in our mass-rearing indicated they were haploid, suggesting that the two-step mechanism of endosymbiont-induced thelytoky is not applicable in this wasp.

Therefore, we investigated the mechanism by which *Wolbachia* induces thelytoky in *M. uniraptor* and propose a model for the sex determining cascade to understand the mechanism by which feminization could be achieved. First, we assessed the frequency of haploid and diploid male offspring after selective removal of *Wolbachia* from *M. uniraptor* females using an increasing concentration of tetracycline. The sex determination genes in *M. uniraptor* share features with those of other hymenopterans, and *N. vitripennis* in particular, in terms of splicing structures and function (Wang, 2021). Based on this knowledge, we next analysed the association of *Wolbachia* presence with both maternal and zygotic expression and splicing of sex determination genes. As several sexual reproduction barriers prevent the establishment of *M. uniraptor* sexual lines, including a reduced ability to store sperm, female mating reluctancy and male infertility (Gottlieb and Zchori-Fein, 2001), we used the sister species *M. raptorellus* to generate triploid females that would asexually produce haploid and diploid offspring, to investigate whether diploidization is sufficient for feminization in this genus.

Our results suggest that in the genus *Muscidifurax*, being diploid is sufficient for female development. We prove that endosymbiont-induced thelytoky can be a one-step process, in which *Wolbachia* only has to induce diploidization which is automatically followed by feminization. Our analysis of the *M. uniraptor* sex determination mechanism is discussed in the light of the different mechanisms that *Wolbachia* can employ to induce thelytoky in haplodiploid insects.

## Material and method

### Insect rearing

*Muscidifurax uniraptor* and *M. raptorellus* were kindly provided to us by Jack Werren and Koppert Biological Systems respectively. The two cultures were continuously reared on *Calliphora spp*. pupae under laboratory conditions of light/dark 16h/8h at 25°C.

### Partial Wolbachia removal from M. uniraptor female

To reduce the *Wolbachia* titre in *M. uniraptor* females, one 10% sucrose control solution and seven different concentrations of tetracycline solutions: 5×10^−2^%, 2.5×10^−2^%, 1.25×10^−2^%, 5×10^−3^%, 2.5×10^−3^%, 1.25×10^−3^% and 5×10^−4^% were prepared by diluting the tetracycline hydrochloride power (Sigma-Aldrich) into 10% sucrose solution. All prepared solutions were filtered by 0.22 μl Millex-GP Syringe Filter (Sigma-Aldrich) and stored at −20°C.

Newly emerged *M. uniraptor* females were collected and transferred to separate glass vials and closed with a cotton plug. A strip of filter paper was placed in each vial with 25 μl of prepared tetracycline solution for 24 hours. Each treatment included 40 individual females. Ten females of each treatment were subsequently offered five *Calliphora* hosts to lay eggs, every 24 hours, for three consecutive days. Parasitized hosts were collected separately and incubated at rearing conditions until offspring emerged. For each mother, the offspring numbers and their sex was recorded. Since offspring emerging from the host presented on the first day were almost all females across all treatments, the offspring that emerged from the second- and third-day parasitized hosts was used for further ploidy analysis. Experiments conducted by Ma et al. (2015) suggest that diploid male offspring often has an intermediate *Wolbachia* titre, which we assumed to have in samples from our low concentration tetracycline treatments. Therefore, the heads from male wasps emerging from the four lowest tetracycline concentration treatments 5×10^−3^%, 2.5×10^−3^%, 1.25×10^−3^% and 5×10^−4^% were collected and stored at −20°C for ploidy check. The remaining 30 mothers from each treatment were dissected in a drop of 1 X PBS solution on glass slides to collect ovaries. Five ovaries were pooled together as one replicate, and six replicates per treatment were collected in 1.5 ml Eppendorf tubes and were fast frozen in liquid nitrogen before storage in – 80°C.

### Offspring ploidy level determination

Flowcytometry was used to identify the ploidy level of offspring from different tetracycline treatments following the protocol adapted from Ma et al. (2015). One frozen wasp head was put into an Eppendorf tube with 0.3 ml ice-cold Galbraith buffer. A plastic pestle was used to crack open the head and homogenize the sample. The homogenate was then poured through a 5 ml cell strainer cap (Stemcell Technologies) to filter out the debris, and was subsequently collected in 96-well plates. Samples were stained with 5 μl Propidium iodide (1.0 mg/ml, Biotium). The total DNA content of each sample was measured by a MACSQuant Analyzer (Miltenyi Biotec) and visualized by a log scale generated fluorescence intensity plot using MACSQuantify™ Software 2.11. Representative figures were generated by gating the raw data in Flowjo v10.6.2 (Becton, Dickinson & Company).

### Collecting embryo stage samples

Newly emerged females were collected in separate glass vials, closed with a cotton plug and fed with either 25 μl of 10% sucrose solution for *Wolbachia*-positive control (*Wol+*) or 25 μl of 0.5% tetracycline in 10% sucrose solution for *Wolbachia*-negative treatment (*Wol-*). Solutions were first provided on strip-type filter papers to females for 12 hours. Then, the filter papers were removed and two hosts were provided per single female to lay eggs within 12 hours. These procedures were applied for two consecutive days prior to embryo egg collection. In order to confirm complete *Wolbachia* removal, the parasitized hosts from these two days were collected and incubated at rearing conditions for offspring sex check. To facilitate the embryo egg collection, an egg-laying chamber was used as described in Wang et al. (2020). Fed females were transferred into the assembled vials in which they could only access the head portion of the fly host. After 24 hours training in the egg-laying chamber with a single host, females were given two hours to lay eggs in the hosts and 30 mins was used for egg collection. For later time point samples, parasitized hosts were first collected after a two-hour egg-laying period and incubated in the rearing condition before the embryo eggs were collected. Eight time windows of 0-2.5h, 2.5-5h, 5-7.5h, 7.5-10h, 10-12.5h, 12.5-15h, 15-17.5h and 17.5-20h were set for embryo collection. A needle with a hook was used for embryo collection to open the shell of the fly pupa and transfer the embryos into a 1.5 ml Eppendorf tube filled with 10 μl of absolute ethanol. More than 40 embryo were pooled as one sample and three replicates were collected per time point of both *Wol+* and *Wol*-treatments. The embryo samples for the *gfp* spike-in experiment were prepared as described before and were collected at 7.5-10h for both *Wol+* and *Wol-* treatment. Each sample contained exactly 35 embryos.

### Maternal provision and zygotic expression of *Mutra* and *Mutra2*

To analyse the maternal provision and zygotic expression of *Mutra* and *Mutra2* alternative splice forms, collected ovaries or embryos were homogenized by plastic pestles and total RNA was extracted using ZR Tissue & Insect RNA MicroPrep™(Zymo Research) following manufacturer’s instructions. For embryo samples, an on-column DNase treatment step (Zymo Research) was added during RNA extraction and 16 μl of DNase/RNase free water was added to each sample to recover the total RNA from the extraction columns. RNA from ovary samples was recovered from the extraction columns by adding 13 μl DNase/RNase free water to each sample. One μl of RNA was used to verify the RNA concentration on a spectrophotometer (DS-11-DeNovix). DNase treatment was carried out on ovary samples using RQ1 RNase-Free DNase (Promega) following manufacture’s protocol. One μg RNA of each ovary sample or 0.5 μg RNA of each embryo sample was synthesized into cDNA by reverse transcriptase PCR (RT-PCR) with standard reaction mix (SensiFAST™ cDNA Synthesis Kit, Bioline) in a thermal cycler (Bio-Rad T100TM Thermal Cycler, Bio-Rad) with 5 minutes priming at 25°C, 15 minutes reverse transcription at 46°C and 15 minutes reverse transcription at 42°C followed by 5 minutes reverse transcriptase inactivation at 85°C. GoTaq® G2 Flexi DNA Polymerase (Promega) was used according to manufacturer’s instructions to prepare the mastermix for PCR (primer concentration: 0.4μM, MgCl^2^ solution: 1.5mM). Target genes *Mutra* and *Mutra2* and internal control gene *MuRP49* were amplified with a standard PCR profile: 3 minutes at 95°C, 32 amplification cycles of 30 seconds at 95°C, 30 seconds at 55°C, 50 seconds at 72°C and a final extension of 5 minutes at 72 °C in a thermal cycler (T100TM Thermal Cycler, Bio-Rad). Primer sequences are listed in Table S1.

Ovary and embryo samples were further analysed with quantitative reverse transcriptase PCR (qPCR) to acquire the relative expression of *Mutra* and *Mutra2*. cDNA that was generated in the RT-PCR was diluted 50 times to be used as qPCR templates and qPCR mastermix was prepared according to SensiFAST™ SYBR® No-ROX Kit manual (primer concentration: 0.4μM, Bioline). *MuRP49* was used as the internal control for ovary samples and *MuEF-1a* was selected as the internal control for embryo samples. *Wolbachia surface protein* (*WSP*) was used for *Wolbachia* quantification. Primer sequences used in qPCR are listed in Table S1. qPCR was carried out using the CFX96TM Real-Time System (Bio-Rad) with Bio-Rad CFX Manager 3.1 Software (Bio-Rad). The standard qPCR profiles were used for all target genes consisting of 95°C for 3 minutes, 40 amplification cycles of 15 seconds at 95°C,15 seconds of 55°C, 30 seconds of 72°C followed by a melting curve check.

### Quantification of Mutra and Mutra2 in embryos using gfp spike-in

To account for the influence of ploidy levels in gene expression difference, we performed the quantification of *Mutra* and *Mutra2* in embryo at 7.5-10h between *Wol+* and *Wol*-treatments using *gfp* spike-in. *Gfp* template was first amplified from the vector pOPINEneo-3C-GFP which was a gift from Ray Owens (Addgene plasmid # 53534; http://n2t.net/addgene: 53534; RRID: Addgene_53534). Subsequently, *gfp* RNA was reverse synthesized using MEGAscript™ T7 Transcription Kit (Thermo Fisher) and purified using MEGAclear™ Transcription Clean-Up Kit (Thermo Fisher) following manufacturer’s instructions. Purified *gfp* RNA was further quantified using Qubit RNA BR Assay Kit (Thermo Fisher) with Qubit 2.0 Fluorometer (Thermo Fisher). Prior to the embryo RNA extraction, 0.03ng *gfp* RNA was added to each sample. Subsequent embryo RNA extraction, cDNA synthesis and qPCR procedures were done as described in the former section. Primers details are provided in Table S1.

### Generating triploid *M. raptorellus* females

To investigate whether female development relies solely on ploidy level in *Muscidifurax* species, triploid females of *M. raptorellus* were created. We used the same parental RNA interference (pRNAi) protocol described in Wang (2021) to maternally silence the *Mrtra* in pupal stage *M. raptorellus* females to first produce diploid males. These diploid males were mated with virgin females to create triploid female offspring. The rearing scheme is illustrated in Figure S1.

Initially, 40 F0 female pupae of *M. raptorellus* were injected with *Mrtra* dsRNA. Once these injected F0 females emerged, we kept them in separate glass vials sealed with cotton plugs. A single male was provided to each F0 female for 24 hours for mating. Thereafter, one fly pupa host was offered to each female and another 24 hours were given for oviposition. Subsequently, the F0 females were freeze-killed and stored in −80°C, and parasitized pupa hosts were incubated in rearing condition until F1 adult emerged. Eight F0 females that produced only male offspring were chosen and their offspring was used for the next step. All F1 males (being either haploid or diploid) from the eight selected F0 females were mated with a virgin female for 24 hours and subsequently freeze-killed and stored in −80°C. Each female that mated with a F1 male was given a single fly pupa host for oviposition for 24 hours, resulting in F2 female offspring that was subsequently collected before eclosion and separated in individual glass vials. These F2 females (containing both triploid and diploid individuals) were further provided with five fly hosts per day for four consecutive days to boost their offspring production. When we observed females in F3, these F3 females were used for further analysis and their male siblings (if present) were freeze-killed and stored in – 80°C. To assess the reproductive ability of females uniparentally produced by triploid mothers, we set them up to produce offspring as virgins and after mating. We collected five F3 virgin female offspring of triploid mothers and provided them with two fly pupa as hosts per female to check their asexual reproductive ability and offspring sex. The remaining F3 females were given 24 hours to mate with males, and two hosts per mated female were offered for 24 hours to check their sexual reproductive ability (including mating) and offspring sex. After oviposition, tested F3 females were freeze-killed and stored in −80°C. All the offspring in each generation that was descendent from same F0 female, F1 male and F2 female was noted and the numbers of male and female offspring of each generation from each mother were counted. After collecting all samples, the heads of the wasps were used to verify the ploidy level in flowcytometry following the same procedures described in former section.

### Data analysis

qPCR data was first imported to LinRegPCR software (LinRegPCR, 2017.1.0.0, HFRC, Amsterdam, The Netherlands) (Ramakers et al., 2003). After baseline correction, the initial number of templates (N0) were calculated based on the average PCR efficiency of each amplicon. Relative expression levels of *NvDsx* in each sample was obtained by dividing the N0 value of *NvDsx* by N0 value of internal or external reference genes. The absolute mass of each transcript in the *gfp* spike experiment is based on the ratio of its N0 and *gfp* N0 and multiplied by 0.03. Then, the exact transcript number is calculated using the formula:

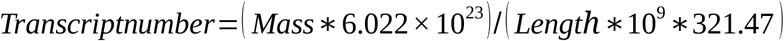

One mole is 6.022×10^23^ molecules; Length – transcript length (*Mutra*: 172bp, *Mutra2*:184bp, *WSP*: 174bp); Mass – Average weight of an RNA base pair (bp) is 321.47 Daltons.

Statistical analysis was performed in R (R core Team, 2018). Ovary qPCR data was analysed using One-way ANOVA and Tukey’s honest significant difference (HSD) as post hoc test or Kruskal-Wallis multiple comparison Dunn-test; embryo different developmental stage and *gfp* spike-in qPCR data were analysed by *t*-test and corrected with false discovery rate (FDR), and Kruskal-Wallis rank sum test. Triploid female offspring data was analysed using Kruskal-Wallis rank sum test and Dunn-test.

## Results

### Reduction of *Wolbachia* titre in *M. uniraptor* female ovaries leads to haploid male offspring production

To investigate thelytokous reproduction induced by *Wolbachia* in our study species *M. uniraptor*, we first asked if diploidization and feminization is decoupled by reducing the amount of *Wolbachia* in females. We expected to find diploid male offspring if *M. uniraptor* follows a two-step feminization model as described in Ma et al. (2015). We reduced the *Wolbachia* titre in a step-wise fashion by feeding increasing concentrations of tetracycline to different groups of freshly emerged female wasps. The results show that we managed to partially remove *Wolbachia* in different treatments and, importantly, that *Wolbachia* titre in female ovaries is inversely correlated with increasing tetracycline concentrations (Figure 1A). A decreased *Wolbachia* titre directly affects the percentage of male offspring which decreased from around 70% in the treatment with the highest tetracycline concentration to almost 0% in the treatment with lowest concentration (Figure 1B). We observed that female offspring always emerged from the fly hosts provided on the first day, regardless of the tetracycline concentration treatment. Therefore, we could not obtain a complete sex reversal even in the treatment with the highest antibiotic concentration. We assume that the penetration of antibiotic was incomplete in the first set of matured eggs, and penetration increased with egg maturation over the next two consecutive days as was shown by Zchori-Fein et al. (2000). In four treatments with lower tetracycline concentrations, *Wolbachia* titre was reduced to intermediate levels in the female ovaries, and after examining offspring ploidy in these treatments, we found no diploid males (Figure 1C, S2C,D).

**Figure 1:**
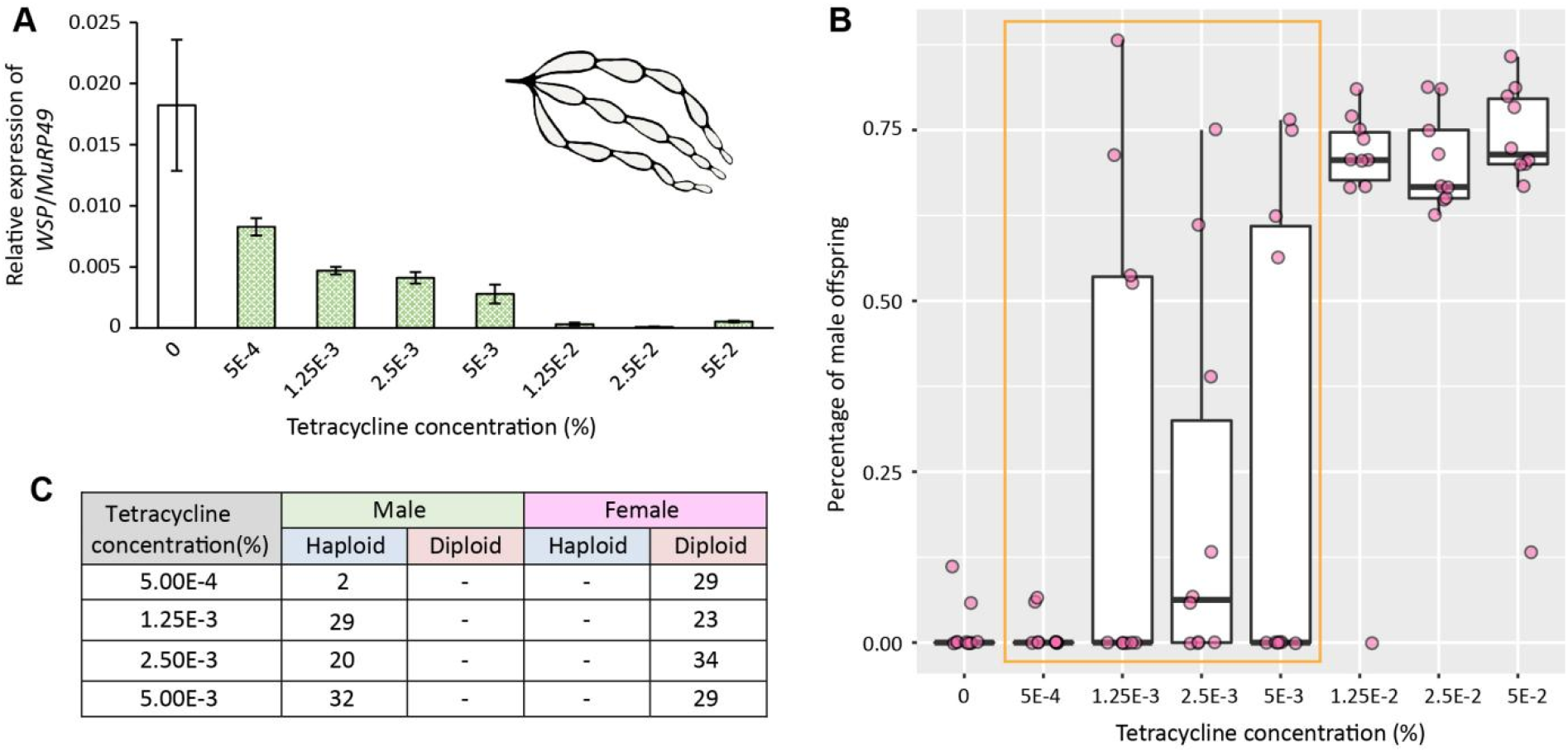
Lower *Wolbachia* titre is inversely correlated with increasing percentage of haploid male offspring in tetracycline treated females. (A) Relative expression of *Wolbachia* gene, *WSP/MuRP49*, in different tetracycline treated female ovaries. The concentration of tetracycline solution of each treatment is indicated on the X-axis. Mean plus standard error is given for each treatment (n=6). (B) Percentage of male offspring from females treated with increasing concentrations of tetracycline. The box-and-whisker plots show median (horizontal line), 25-75 % quartiles (box) and maximum/minimum range (whiskers). The pink data points represent the individual sample points (n=10). The yellow box indicates the treatments that are used for ploidy level confirmation. The concentration of tetracycline solution of each treatment is indicated on the X-axis. (C) Ploidy level of the offspring produced by female treated with different concentrations of tetracycline.

### *Mutra* and *Mutra2* expression is not affected by *Wolbachia* density in the ovaries

Previous data confirmed that the maternal provision of both *Mutra* and *Mutra2* plays an essential role in female development (Wang, 2021). Silencing *Mutra* or *Mutra2* in female *M. uniraptor*, results in 100% male offspring regardless of the presence of *Wolbachia*. We hypothesized that *Wolbachia* is involved in the regulation of maternal provision of *Mutra* and *Mutra2* transcripts to the eggs. Therefore, we performed both RT-PCR and qPCR to study their expression pattern and expression level in tetracycline-treated female ovaries. The expression level of *Mutra* varied among treatments with increasing dose of tetracycline (Figure 2A). We conclude that there is no correlation between *Mutra* expression and *Wolbachia* titre (Figure 2A). *Mutra2* expression on the other hand, significantly increased in the 1.25×10^−3^% tetracycline-treated group compared to the non-treated group (*P*=0.028, Figure 2B). However, in the treatment groups with increasing dose of tetracycline, *Mutra2* expression decreased to levels comparable to the non-treated group (Figure 2B). We cannot explain the variable levels of *Mutra2* in response to tetracycline treatment of the females and suggest that observed differences in *Mutra2* expression are due to biological variation and are not related to *Wolbachia* titre (Figure 2B).

**Figure 2:**
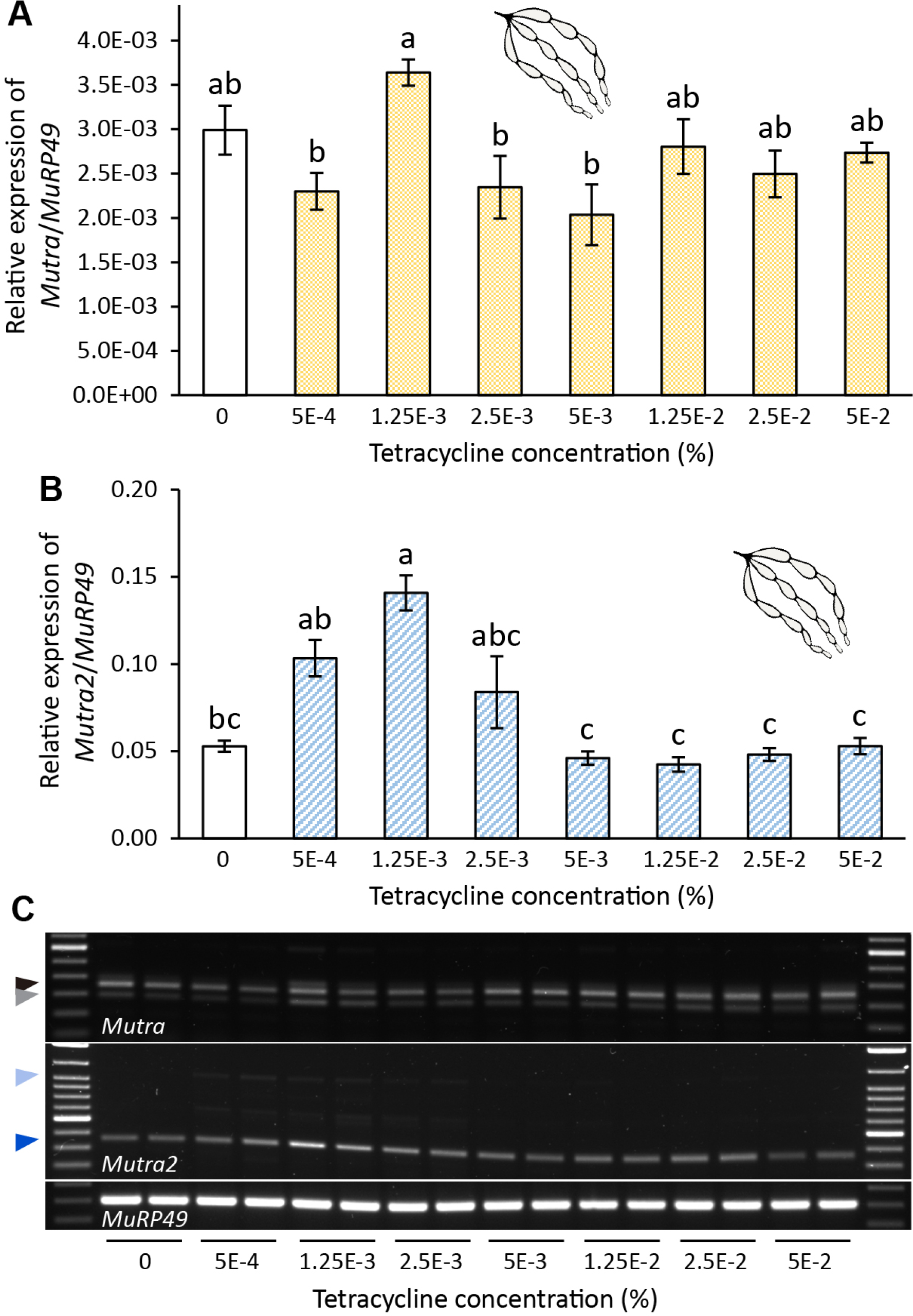
*M. uniraptor transformer* (*Mutra*) and *M. uniraptor transformer-2* (*Mutra2*) expression and splicing patterns in the ovaries of females treated with different concentrations of tetracycline. Relative expression of **(A)** *Mutra/MuRP49* and **(B)** *Mutra2/MuRP49* in ovaries of *M. uniraptor* females treated with increasing dose of tetracycline concentration (x-axis represents the percentage of tetracycline solution). Error bars represent standard error (SE) (n=6). Statistics was performed with One-way ANOVA and Tukey’s HSD for A, with Kruskal-Wallis multiple comparison Dunn-test for B. Letters refer to statistically significant differences with *P*<0.05. **(C)** Gel electrophoresis of RT-PCR products of *Mutra* and *Mutra2* splice variants and *MuRP49* fragments in ovaries of *M. uniraptor* females treated with increasing tetracycline concentration indicated under the gels. Black and grey arrows refer to non-sex-specific (NSS) and female-sex specific (F) *Mutra*, dark blue and light blue arrows refer to more abundant non-sex specific splicing *Mutra2^NSS1^* and *Mutra2^NSS2^*. *MuRP49* serves as internal control and 100bp ladder (Thermo Fisher) indicates fragment size. Fragments are visualized on 1.5% TAE agarose gel stained with Midori Green (NIPPON Genetics).

Next, we hypothesized that *Wolbachia* is involved in the sex-specific splicing of *Mutra* and *Mutra2* transcripts that are maternally provide to the eggs which would result in a shift from female-specific splicing to male-specific splicing. Currently, three *Mutra* splice variants have been identified in *M. uniraptor*, including a male-specific *Mutra^M^*, a female-specific *Mutra^F^*, and a non-sex-specific *Mutra^NSS^* (Wang, 2021). Both *Mutra^F^* and *Mutra^NSS^* were found to be expressed in the female ovary (Figure 2C) while only *Mutra^F^* was present in the rest of the female body (Figure S3). The splice variant *Mutra^F^* and *Mutra^NSS^* are present in all treatment groups (Figure 2C), and a relatively higher expression level of *Mutra^NSS^* compared to *Mutra^F^* was observed. However, expression of both splice variants is similar across different treatment groups (Figure 2C). The non-sex specific *Mutra2^NSS1^* is predominantly expressed in all treatment groups (Figure 2C). *Mutra2^NSS2^* splice form was found to be highly abundant in whole body tissues in different developmental stages of female (Wang, 2021) but is barely expressed in ovaries (Figure 2C). Concludingly, we observed no shift in *Mutra* or *Mutra2* splicing patterns among treatments with increasing dose of tetracycline (Figure 2C), and we determine that *Wolbachia* is not involved in sex-specific splicing of *Mutra* or *Mutra2* in female ovaries.

### *Wolbachia* affects zygotic expression of *Mutra* but not *Mutra2* during early embryogenesis

*Wolbachia* infection does not affect expression levels or splicing patterns of maternal provision of *Mutra* and *Mutra2*, still, a reduction in *Wolbachia* titre results in a shift of offspring sex to males. Therefore, we hypothesized that *Wolbachia* manipulates sex determination during early zygotic development by changing zygotic expression of sex determination genes. We analysed the zygotic expression profiles of *Mutra* and *Mutra2* in embryo samples from both tetracycline-treated (*Wol*-) and non-treated (*Wol*+) females that were collected in eight time windows from zero to 20 hours post oviposition (hpo). We first confirmed that *Wol*-zygotes have significantly reduced *Wolbachia-*infection throughout zygotic development, showing the success of antibiotic treatment (Figure 3A). *Mutra* expression increased from zero to 10 hpo and showed peak expression at 7.5-10 hpo (Figure 3B). No significant difference of *Mutra* expression between *Wol*-and *Wol*+ treatments was found in 0-2.5 hpo, corroborating that *Wolbachia* does not affect maternal provision of *Mutra*. From 2.5-7.5 hpo onwards there is no difference in *Mutra* expression between *Wol*+ and *Wol*-. After the peak, *Mutra* expression remains consistent from 10 hpo onward with no significant differences in both treatments, although overall a slightly higher expression of *Mutra* was observed in *Wol*-treatment (Figure 3B). Next, we determined *Mutra* sex-specific splicing during zygotic development in *Wol*- and *Wol*+ embryos (Figure 3D). From 0-5 hpo, the maternal provision of *Mutra^NSS^ and Mutra^F^* is visible in both *Wol*- and *Wol*+ treatments (Figure 3D). The zygotic expression of *Mutra^F^* increases visibly from 5-10 hpo in *Wol*+ embryos and only from 7.5-10 hpo in *Wol*-embryos except for one sample that increases visibly from 5-10 hpo in *Wol*-embryos. Overall, *Mutra* expression starts earlier and appears stronger in *Wol*+ embryos compared to *Wol*-embryos in the 5-10 hpo developmental window. From 10 hpo onwards a clear shift in sex-specific splicing from *Mutra^F^* to *Mutra^M^* and *Mutra^NSS^* can be observed in *Wol*-embryos, whereas *Wol*+ embryos maintain *Mutra^F^* splicing (Figure 3D). Therefore, we conclude that in *Wol*-embryos male development initiates from 10 hpo, but the effect of *Wolbachia* infection on the zygotic *Mutra* expression levels are not conclusive from this experiment.

**Figure 3:**
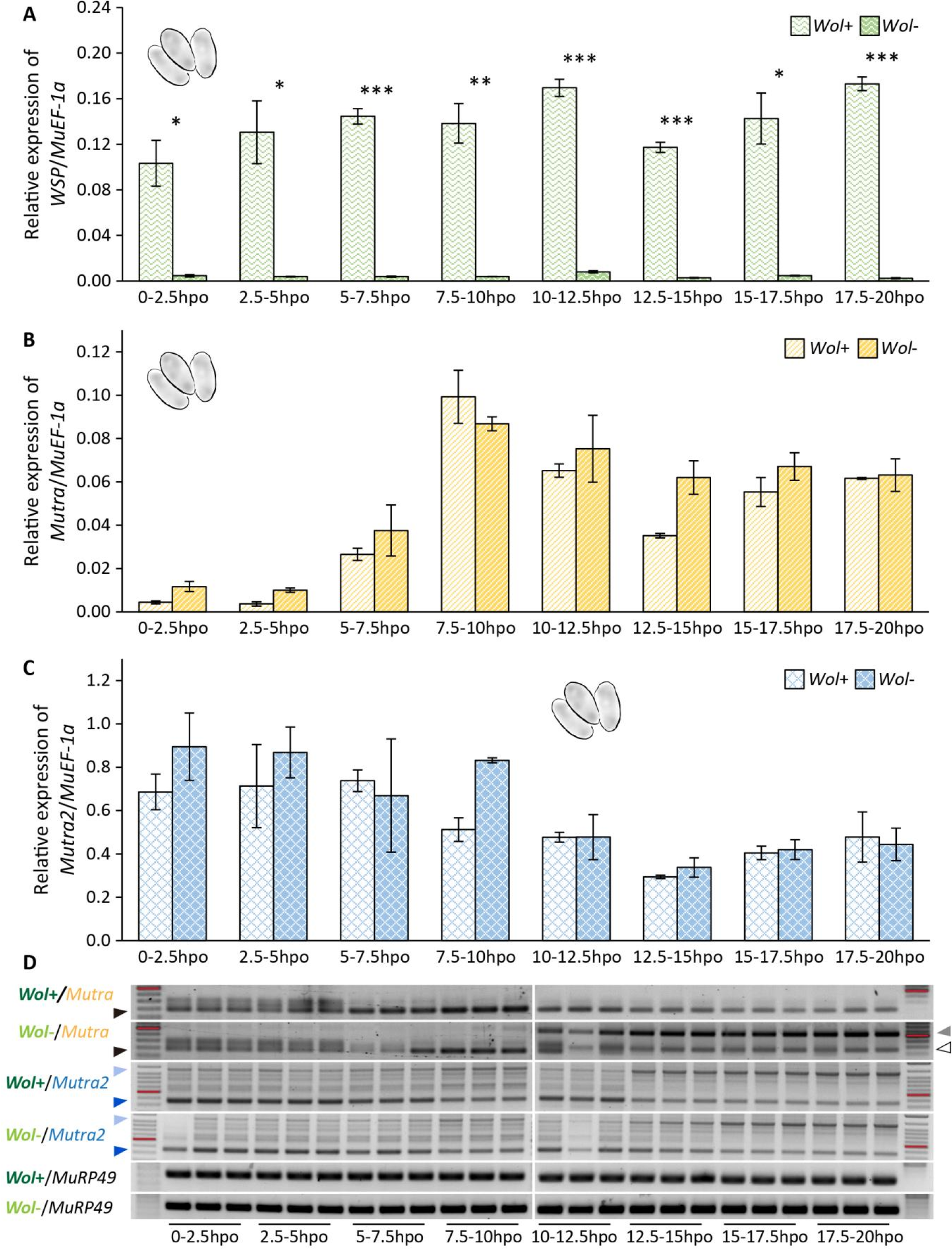
Expression of *M. uniraptor transformer (Mutra)* and *M. uniraptor transformer-2* (*Mutra2*) in early embryogenesis of *Wolbachia* positive (*Wol+*) and *Wolbachia* negative (*Wol-*) embryos. Figures show the relative expression of **(A)** *Wolbachia* gene, *WSP/MuEF-1a*, **(B)** *Mutra/MuEF-1a* and **(C)** *Mutra2/MuEF-1a* in embryo samples from both tetracycline-treated (*Wol*-) and non-treated (*Wol*+) females at eight developmental time windows from zero to 20 hours post oviposition (hpo). Error bars are standard errors with n=3. Statistics was performed with *t*-test and corrected with false discovery rate (FDR) for A, B and C. Asterisks were assigned for significant differences: * = *P*<0.05, ** = *P*<0.01 and *** = *P*<0.001. **(D)** Gel electrophoresis of RT-PCR products of *Mutra* and *Mutra2* splice variants and *MuRP49* fragments in *Wol-* embryos and *Wol*+ embryos at eight developmental time windows from zero to 20 hours post oviposition (hpo). Black, grey, white, light blue and dark blue arrows indicate *Mutra^F^*, *Mutra^M^*, *Mutra^NSS^*, *Mutra2^NSS2^* and *Mutra2^NSS1^* respectively. MuRP49 serves as internal control and 100bp ladder (Thermo Fisher) indicates fragment size (600bp fragments in ladders are marked with red lines). Fragments are visualized on 1.5% TAE agarose gel stained with Midori Green (NIPPON Genetics).

*Mutra2* is also maternally provided to the eggs (0-2.5 hpo) and shows no difference in expression in both *Wol*+ and *Wol*-embryos from all time windows. *Mutra2* expression is approximately 10-fold higher than *Mutra* expression at 7.5-10 hpo, and remains relatively high until 10 hpo after which it gradually decreases (Figure 3C). Next, we determined *Mutra2* sex-specific splicing during early embryogenesis and observed that in both *Wol*+ and *Wol*-treatments splice variants remain unchanged (Figure 3D). Most abundant splice variants are *Mutra2^NSS1^* and *Mutra2^NSS2^*, but also the less abundant splice variants *Mutra2^NSS3^* and *Mutra2^NSS4^* are visible (Figure 3D). We observed that *Mutra2^NSS1^* is expressed predominantly from 0-12.5 hpo in both treatments and decreases in expression from 12.5 hpo onwards, while *Mutra2^NSS2^* expression increases from 12.5 hpo onwards in both treatments (Figure 3D). We conclude that *Wolbachia* does not affect *Mutra2* splicing and expression.

We could not definitively conclude whether or not *Wolbachia* affects *Mutra* expression using relative quantification, but we asked if a diploid individual would have a double dose of *Mutra*, caused by bi-allelic expression, when compared to *Mutra2* expression. Therefore we performed an absolute quantification of *Mutra* and *Mutra2* expression in *Wol*- and *Wol*+ embryos. We collected an equal number of embryos per sample for both treatments, and added a predefined amount of an exogenous synthesized *gfp* RNA (spike RNA) as external standard in every sample before RNA extraction. In this way we could calculate the exact initial RNA input of different samples and correct this against the amount of *gfp* spike RNA for technical variation. As relative *Mutra* expression increases rapidly from 7.5 hpo and the shift of female-specific splicing to male-specific splicing occurs from 10 hpo, we chose to sample at the 7.5-10 hpo time window.

We first confirmed that also in this experiment *Wol*-embryos have a significant reduction of *Wolbachia* (*P*=0.006, Figure 4A). Next, we found that the *Mutra* expression in *Wol*+ embryos is more than two-fold compared to the *Mutra* expression in *Wol-* embryos (*P*=0.003, Figure 4B). As expected, we did not find a significant difference in *Mutra2* expression between *Wol*- and *Wol*+ treatments (*P*=0.535, Figure 4C). These results show that a *Wolbachia*-infected diploid individual has a two-fold higher *Mutra* expression than an uninfected haploid individual, suggesting that either *Wolbachia* regulates *Mutra* expression or *Mutra* has bi-allelic expression compared to *Mutra2* and possibly other zygotic genes. This hints towards a sex-determination model in which diploidization results in enough *Mutra* expression to maintain the positive autoregulatory loop of female-specific *Mutra* splicing as shown for other Hymenoptera.

**Figure 4:**
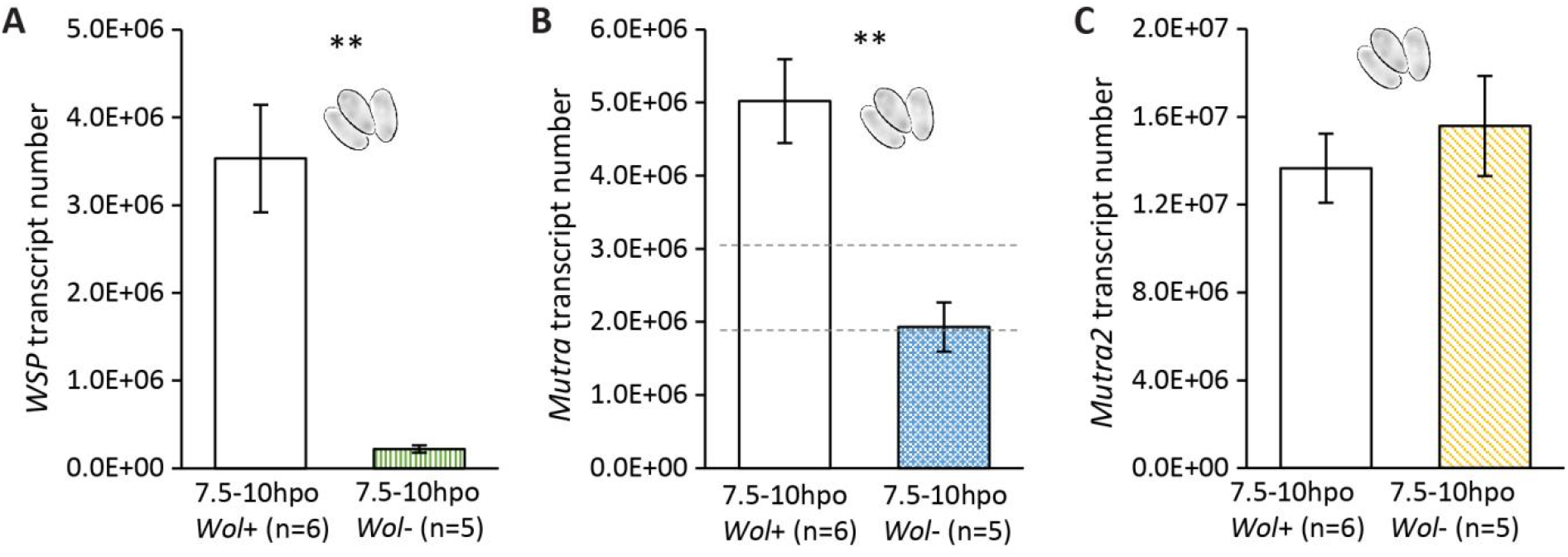
Expression of *WSP*, *Mutra* and *Mutra2* in *Wolbachia* positive (*Wol+*) and *Wolbachia* negative (*Wol-*) embryos at 7.5-10 hours post oviposition (hpo). Figures show **(A)** *WSP*, **(B)** *Mutra* and **(C)** *Mutra2* transcript number in embryo samples from both tetracycline-treated (*Wol-*) and non-treated (*Wol+*) females. RNA spike is *gfp*. Error bars represent standard errors. Dashed lines in (B) represent the upper and lower range of mono-allelic *Mutra* expression in *Wol+* control embryos. Statistics was performed with Kruskal-Wallis rank sum test for A, with One-way ANOVA for B and C. Asterisks represent significant difference: ** = P<0.01.

### Ploidy level determines the sex in *M. raptorellus* and possibly *M. uniraptor*

In *M. uniraptor*, *Wolbachia* induces diploidy restoration of unfertilized eggs after completion of the first mitosis directly after egg laying (Gottlieb et al., 2002). We hypothesized that in *M. uniraptor*, *Wolbachia*-induced diploidization is sufficient for feminization and *Wolbachia*-induced thelytoky is a one-step mechanism. In *N. vitripennis* it has been shown that triploid females produce diploid and haploid offspring when virgin (Beukeboom and Kamping, 2006), and triploid females can be artificially created by mating a diploid female to a diploid male (Leung et al., 2019). We show have shown before that parental RNAi of *Mutra* or *Mrtra* results in a shift of female development into male development, thereby creating functional diploid males (Wang, 2021). In order to prove our hypothesis, we used *M. raptorellus*, a sexually reproducing sister species of *M. uniraptor* to first generate diploid males by maternally silencing *Mrtra* in females and then mate these diploid males with normal diploid *M. raptorellus* females to obtain triploid females (Figure S1). After successfully silencing *Mrtra* in F0 females (Figure 5A), these females were mated to normal haploid males to produce diploid male F1 offspring (Figure S1). These diploid males were subsequently mated to normal diploid females to produce triploid female offspring and haploid male offspring in the F2 (Figure S1).

**Figure 5:**
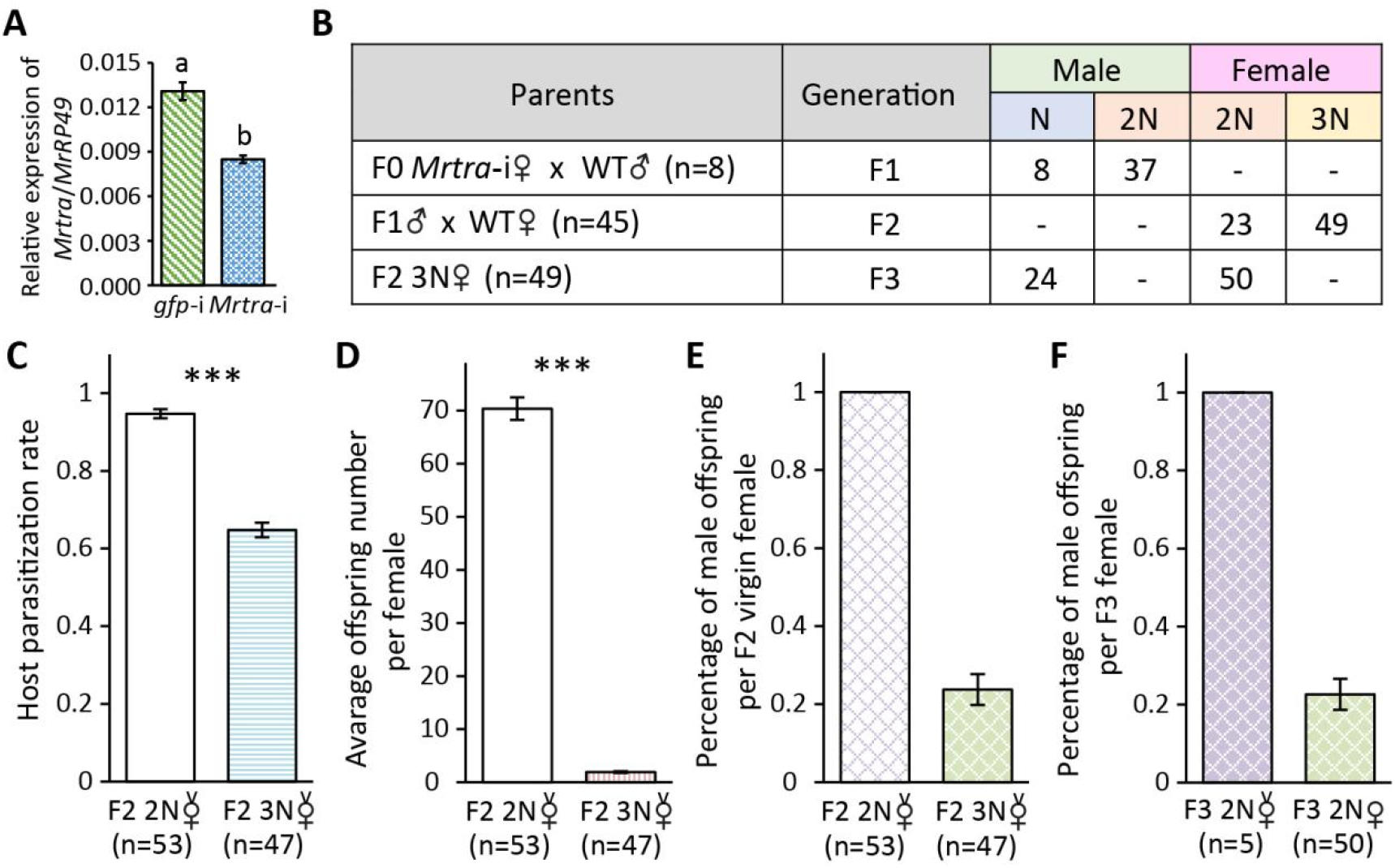
Triploid *M. raptorellus* females produce impaternate (uniparental reproduction by mother) diploid females but have low fecundity. **(A)** Relative expression of *Mrtra/MuRP49* in *Mrtra* silenced (*Mrtra*-i) females and *gfp/MuRP49* in *gfp* silenced (*gfp*-i) females. **(B)** Table showing the number of parental crosses to generate polyploid offspring, the generation number, and the amount of haploid, diploid and triploid offspring and their sexes. Number of parental crosses in each generation is indicated in the column “Parents” with “n”. “N”, “2N” and “3N” refers to haploid, diploid and triploid male or female offspring. **(C)** Host parasitization rate of F2 diploid and triploid females. **(D)** Average offspring number of F2 diploid and triploid females. **(E)** Proportion of male offspring produced by F2 diploid and triploid virgin females. **(F)** Proportion of male offspring produced by F3 diploid virgin and mated females. “2N” and “3N” in C, D, E and F refers to diploid and triploid females. “v” on top of the female symbols represents virgin females. Error bars are standard error of A (n=4), C, D, E and F. Statistics was performed with *t*-test for A, with Kruskal-Wallis rank sum test for C, D. Asterisks represent significant difference: *** = *P*<0.001.

Flowcytometry analysis confirms that we obtained 37 diploid males in the F1 offspring and 49 triploid females in the F2 offspring (Figure 5B, Figure S2H,J). These F2 triploid females were set up as virgin and parthenogenetically produced male F3 offspring and female F3 offspring (Figure 5B, Figure S1), which were confirmed to be haploid and diploid respectively (Figure S2K,L). Compared to the F2 diploid females, these F2 triploid females parasitized significantly fewer fly hosts, and offspring likely suffered from aneuploidy, resulting in extremely low average offspring numbers (Figure 5C,D). However, despite the low fecundity of F2 triploid females, they produced a high proportion of impaternate (uniparental reproduction by mother) female offspring by thelytokous reproduction (Figure 5E). The impaternate F3 diploid females were tested for asexual reproduction, and were mated to normal haploid males to confirm their ability to mate and produce offspring. Result shows that impaternate F3 diploid females parthenogenetically produced only F4 male offspring and after mating produced F4 male and female offspring in a ratio similar to F3 triploid females (Figure 5F). We conclude that in *M. raptorellus* and, by extension, in *M. uniraptor*, being diploid is sufficient for female development.

## Discussion

In this research, we investigated the possible regulatory mechanism for *Wolbachia* to induce thelytoky in *M. uniraptor*. As spontaneous diploid males are reported in several wasp species infected with PI-*Wolbachia*, it was assumed that *Wolbachia*-induced thelytoky would take two steps: diploidy restoration and feminization. We asked if *Wolbachia*-induced thelytoky in *M. uniraptor* also requires two steps. We gradually lowered the *Wolbachia* titre in *M. uniraptor* female ovaries by oral feeding increasing concentrations of tetracycline solution, and observed a proportional increase in the male offspring from *Wolbachia-*reduced females. This result confirms the observation by Zchori-Fein et al. (2000) that female development requires a sufficient amount of *Wolbachia* in the ovaries. Using flowcytometry, we verified that even the lower concentrations of tetracycline treatments led to haploid male offspring development, indicating that in *M. uniraptor* separate diploidization and feminization steps are not needed for thelytokous reproduction. Our result is different from *Asobara japonica* and *Trichogramma kaykai* studies that show that *Wolbachia* uses a two-step mechanism to achieve thelytoky (Ma et al., 2015; Tulgetske, 2010). In addition, this sex determination mechanism supports a long-standing model in Hymenoptera in which ploidy itself determines sex.

Further studies on the sex determination genes *tra* and *tra2* in *Wol*- and *Wol*+ female ovaries suggests that neither the expression levels nor the expression patterns of these genes are correlated with the *Wolbachia* titre, indicating that *Wolbachia* does not manipulate the level of *Mutra* and *Mutra2* maternal provision. By comparing the relative expression of *Mutra* and *Mutra2* between *Wol*- and *Wol*+ treatments during early embryo development, we observed comparable expression level of *Mutra* and *Mutra2* from both *Wol*- and *Wol*+ embryos during eight developmental time windows. Strikingly, in both *Wol*- and *Wol*+ the expression of *Mutra* peaked at 7.5-10 hpo, but the splicing patterns of *Mutra* displayed a clear shift from *Mutra^F^* to *Mutra^NSS^* and *Mutra^M^* from 10 hpo onwards only in *Wol*-treatment and not in *Wol*+ treatment. This suggests that the expression of *Mutra^F^* cannot be maintained from 10 hpo onwards without *Wolbachia*. Absolute quantification of transcript numbers of *Mutra* and *Mutra2* at 7.5-10 hpo showed that *Mutra* expression increased significantly only in *Wol*+ embryos, while no significant changes in the *Mutra2* expression were detected in *Wol*+ embryos compared to *Wol*-embryos. Intriguingly, the reduction of *Mutra* in *Wol*-treatment is within the range of half the *Mutra* expression in *Wol*+ treatment, indicating it could due to the bi-allelic expression of *Mutra* in diploid embryos. This sudden increase of *Mutra* expression at 7.5-10 hpo resembles the more than ten-fold higher expression during peak expression in fertilized embryo of *N. vitripennis* compared to unfertilized embryos (Verhulst et al., 2010a; Zou et al., 2020). Assuming the *tra* auto-regulatory loop is well conserved in *Muscidifurax* as has been shown in almost all identified insect species containing *tra* (Verhulst et al., 2010b), the threshold for the number of *tra* transcripts needed to initiate and maintain the auto-regulatory loop is unclear and could vary among species.

By silencing *Mrtra* in *M. raptorellus* females with subsequent mating, we artificially created diploid males in *M. raptorellus*. Mating these diploid males to normal diploid females generated triploid females and these triploid females in our experiments could parthenogenetically reproduce diploid female and haploid male offspring. This further strengthens the idea that diploidization is the only requirement for female development in *Muscidifurax*. *Wolbachia* would then achieve diploidy restoration in *M. uniraptor* by the end of the first mitosis as shown by Gottlieb et al. (2002), and it would not need to take control of the sex determination system.

One remaining question for our proposed *Muscidifurax* sex determination model is whether there is dosage compensation. Dosage compensation has first been observed in *Drosophila*, in which X-chromosome-linked genes, although having a double copy in females (XX) and one copy in males (XY), show equal expression levels in both sexes. Further research discovered that the number of X chromosomes controls the X-chromosome-located *Sex lethal* (*Sxl*) expression in females (Erickson and Quintero, 2007), which encodes a female-specific RNA-binding protein that not only initiates the female sex determination cascade, but also suppresses the dosage compensation machinery (Kelley et al., 1997). This transcriptionally regulated dosage compensation is commonly found in species with sex chromosomes but receives less attention in haplodiploid species without sex chromosomes. In Hymenoptera, dosage compensation was suggested by Rasch et al. (1977) to account for the ploidy difference between sexes. Interestingly, they found that in *N. vitripennis* haploid embryos have indeed half the amount of DNA in the nuclei compared to diploid embryos, but in adult somatic cells an equivalent amount of DNA per cell is found in both males and females. This dosage compensation therefore is thought to be realized by an endoreplication cycle, in which DNA content is increased without cell division (Edgar and Orr-Weaver, 2001).

Rasch et al. (1977) postulated that dosage compensation by endoreplication possibly takes place several hours before the first larval instar. If this is the case in *Muscidifurax*, the observed two-fold higher expression of *tra* at 7.5-10 hpo is only possible in diploid but not haploid embryo, which is in accordance with our one-step-diploidization-causes-feminization model. Furthermore, if increased zygotic *tra* expression is only exhibited through doubled genomic DNA content, we would expect to see that the expression of *tra* relative to reference gene expression will remain unchanged in later developmental time windows, which seems the case based on our embryo qPCR and RT-PCR results. However, a more precise quantification method would be required for further justification. Given the fact that we only observed an increase of *tra* expression but not *tra2* expression in *gfp*-spiked *Wol*+ embryos, it means that there must be a transcriptional level dosage compensation mechanism to suppress *tra2* expression and activate (or release) *tra* expression in diploid embryos. *Sxl* in *Drosophila* and *femaleless* in *Anopheles gambiae* are X-linked genes that are both involved in sex determination and the transcriptional dosage compensation (Kelley et al., 1997; Krzywinska et al., 2021). If there is an upstream regulator in *Muscidifurax* controlling both sex determination and the transcriptional dosage compensation process, it should be able to specifically increase the zygotic *tra* expression in relation to the ploidy level of the embryo. Another possibility is the involvement of *tra* itself in transcriptional dosage compensation. At the moment we severely lack knowledge in this regard and we have no enough evidence to support either theory, more research is required to fully understand the initial sex determination signal in *Muscidifurax*.

Even if the exact sex determination mechanism remains elusive, our result suggests that the sex determination mechanism in *M. raptorellus* and possibly in *M. uniraptor* are different from the only two currently known sex determination mechanisms in Hymenoptera: maternal effect genomic imprinting sex determination (MEGISD) in *N. vitripennis* (Verhulst et al., 2010a; Verhulst et al., 2013; Zou et al., 2020) and the complementary sex determination (CSD) in *Apis mellifera* (Beye et al., 2003; Gempe et al., 2009; Hasselmann et al., 2008). MEGISD requires an active copy of *wom* provided by the male sperm during fertilization to direct zygotic *tra* expression resulting in female development (Zou et al., 2020). Our result rejects the MEGISD system for *Muscidifurax* by showing that female development in *M. raptorellus* can be independent of paternal input. On the other hand, CSD, in which a homozygous *csd* locus leads to male development (Beye et al., 2003; Gempe et al., 2009) is also not compatible with homozygous parthenogenesis production of *Muscidifurax* females (Heimpel and de Boer, 2008). This again demonstrates the diversity of sex determination mechanisms in insects that is awaited our exploration.

## Acknowledgements

We would like to thank Elzemiek Geuverink, Louis van de Zande, Leo Beukeboom and the WUR inSEXdet team for always being there to discuss sex determination cascades and manipulating endosymbionts. We thank Marcel Dicke for his helpful comments on earlier draft versions; Julia Ruiz Capella, Lina Ojeda Prieto, Roxane Snijders and Sander Koene for their pioneering work in determining the proper concentrations of tetracycline used for oral feeding of *M. uniraptor*; Hans Smid for assisting us with general wasp physiological questions; and Min Xu for helping collect and prepare different wasp samples.

---

**Table S1.**
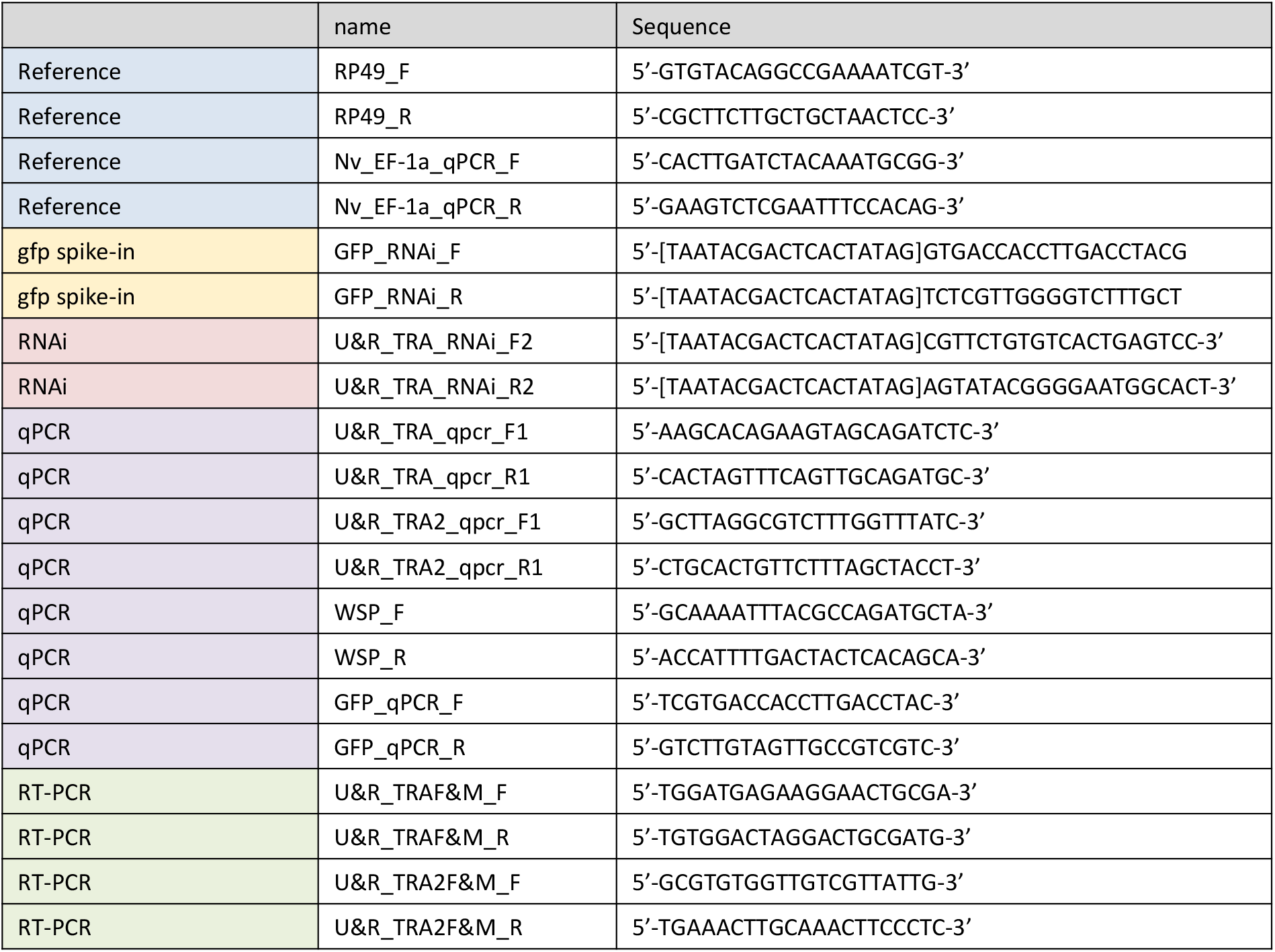
Primers used in this research. “[]” T7 promoter sequence that is recommended by the manufacturer.

**Figure S1:**
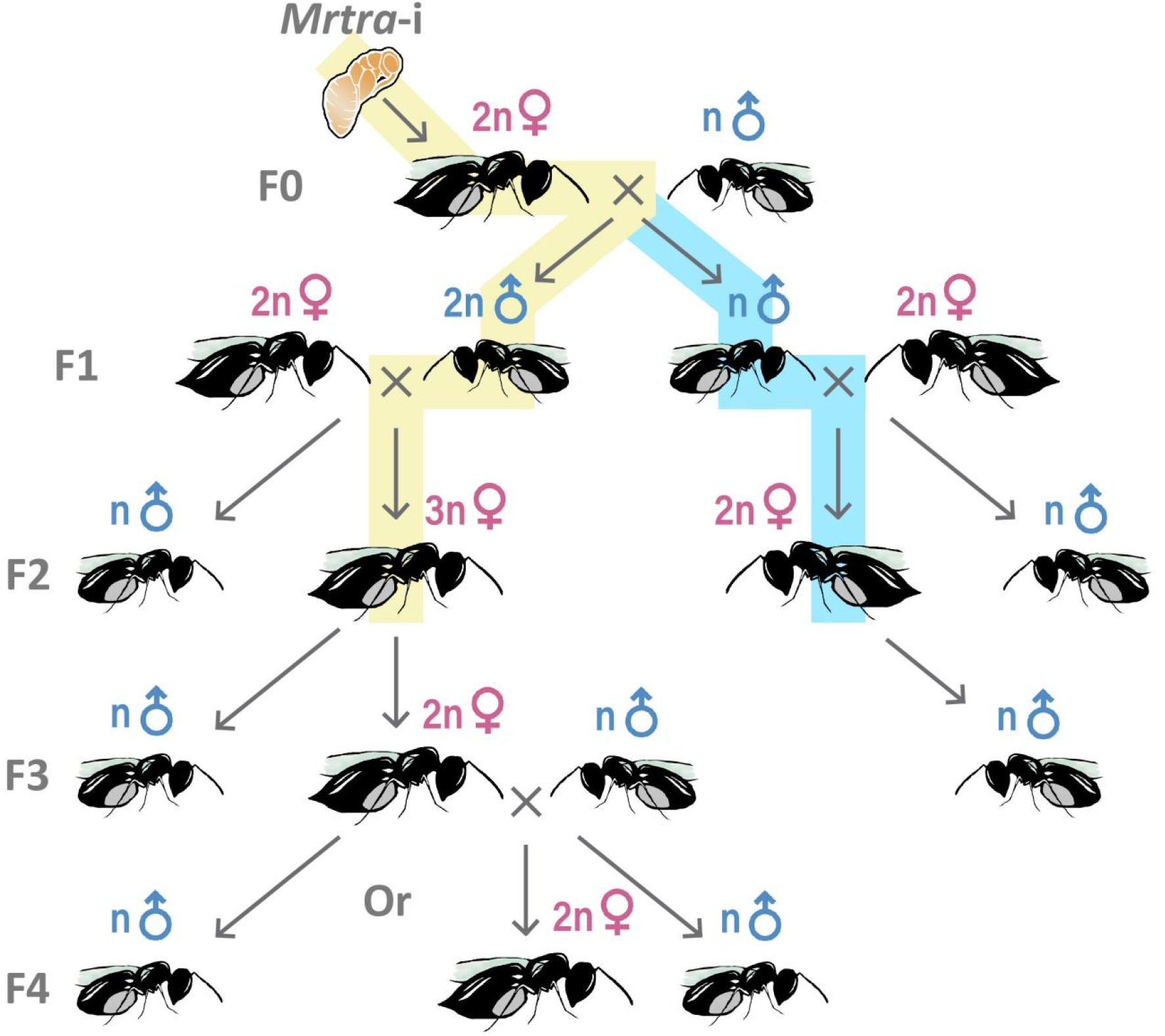
Schematic of the triploid *M. raptorellus* females creation and the resulting offspring sex and ploidy. The generation F0 as indicated on the left of the figure refers to the females treated with *Mrtra* dsRNA (*Mrtra*-i) in the white pupal stage. After emergence they were mated to haploid (n) males and produced haploid and diploid (2n) males (F1). These F1 males were mated to normal diploid females, resulting in the production of F2 triploid (3n) females and haploid males (yellow line, diploid father), or the production of diploid females and haploid males (blue line, haploid father). The F2 triploid females were setup as virgin and produced male and female F3 offspring (yellow line); the F2 diploid female was setup as virgin and produced only male F3 offspring. The impaternate (uniparental reproduction by mother) diploid F3 females were mated to normal haploid males and produced male and female (F4) offspring.

**Figure S2:**
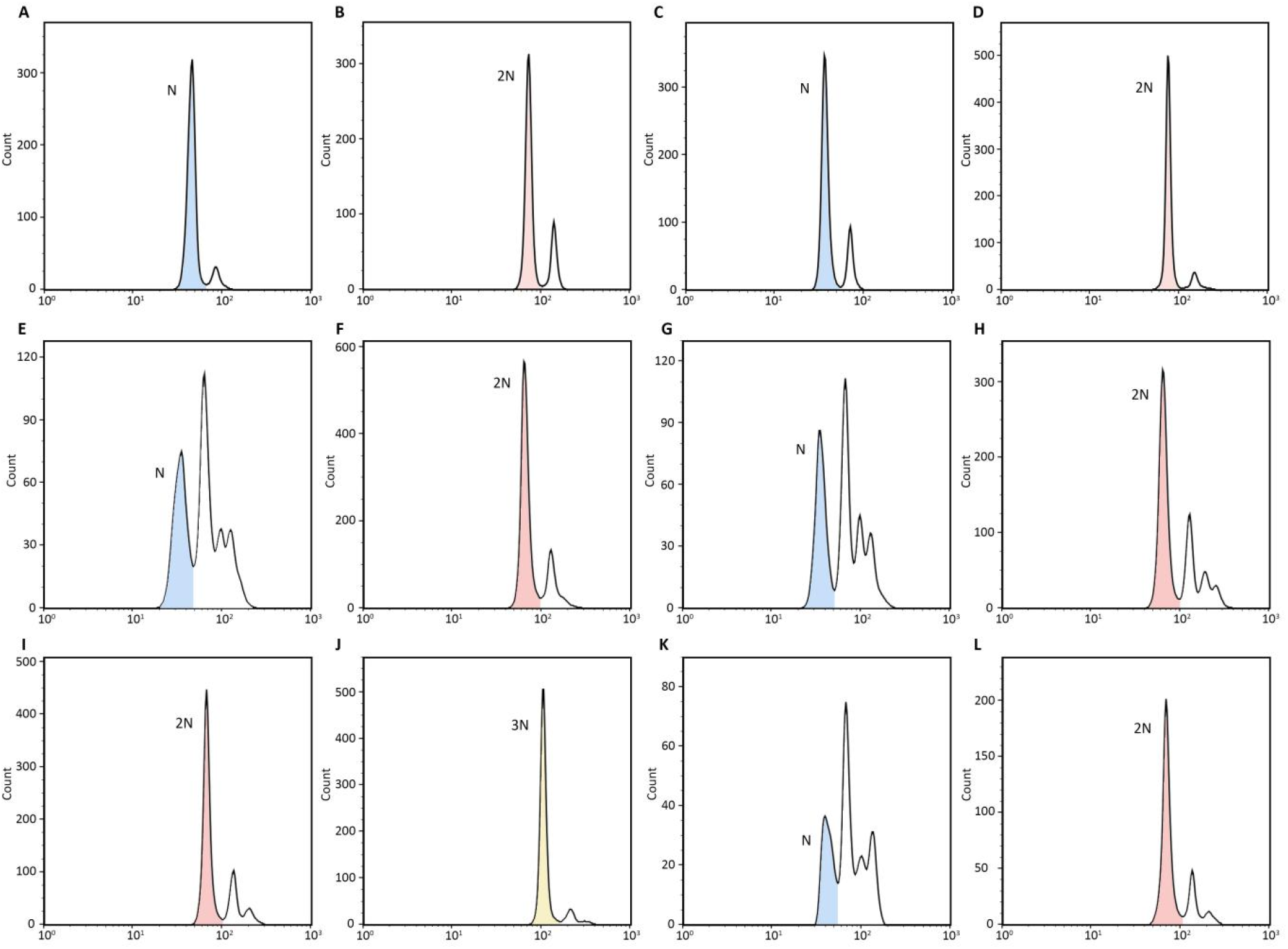
Representative plots depict the ploidy level of wild-type (WT) and offspring male and female from different treatments or generations of *M. uniraptor* and *M. raptorellus*. Peaks that are marked with colour in the figures represent the accumulation of the nucleus in ploidy plots of **(A)** WT *M. uniraptor* male, **(B)** WT *M. uniraptor* female, **(C)** male offspring of tetracycline treated *M. uniraptor* female, **(D)** female offspring of tetracycline treated *M. uniraptor* female, **(E)** WT *M. raptorellus* male, **(F)** WT *M. raptorellus* female, **(G)** *M. raptorellus* haploid F1 male, **(H)** *M. raptorellus* diploid F1 male, **(I)** *M. raptorellus* diploid F2 female, **(J)** *M. raptorellus* triploid F2 female, **(K)** *M. raptorellus* haploid F3 male and **(L)** *M. raptorellus* diploid F3 female. Y axis refers to the number of the nuclei counts. ‘N’, ‘2N’ and ‘3N’ indicate haploid, diploid and triploid represented peaks.

**Figure S3:**
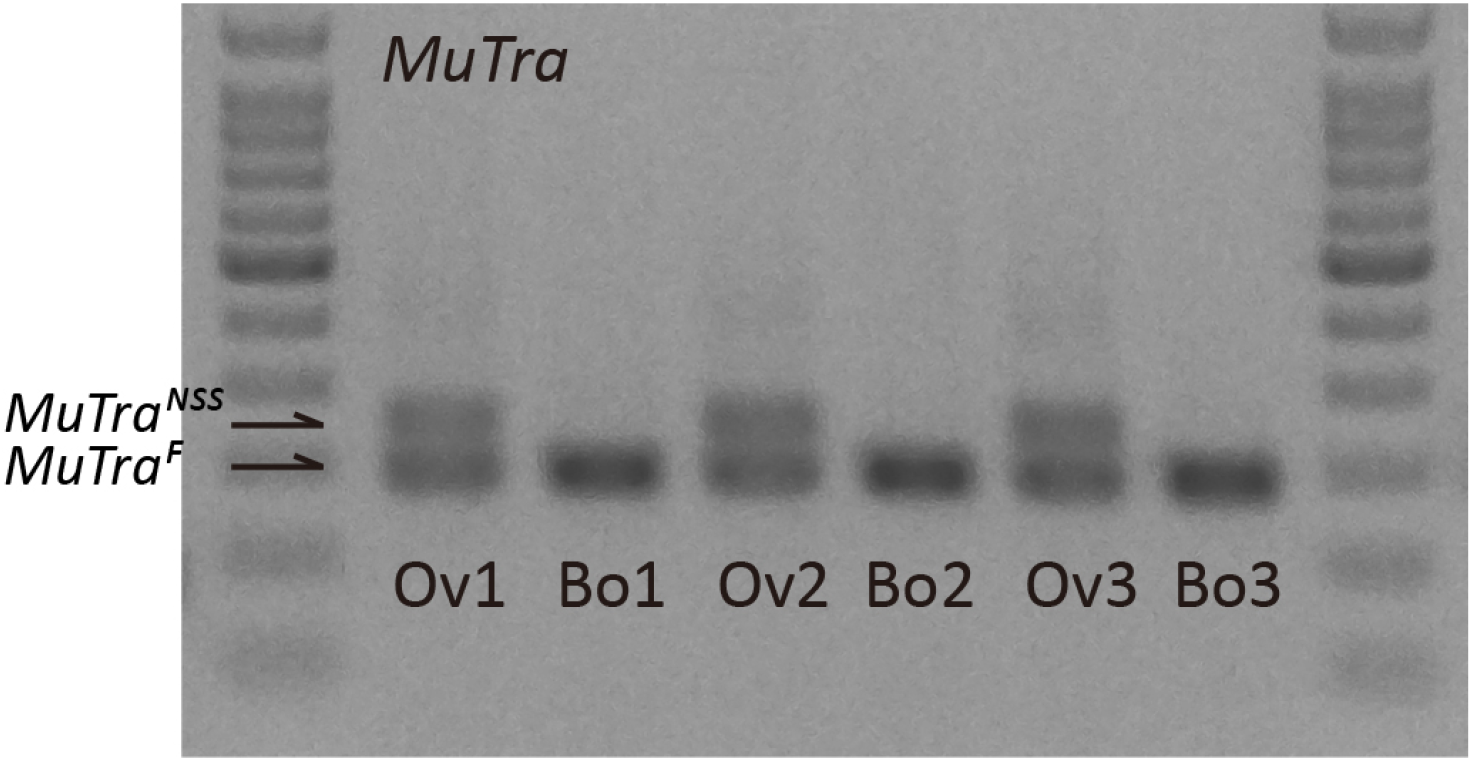
Expression of *Mutra* splicing variants in ovary and body samples of *M. uniraptor*. Female sex-specific (F) and non-sex-specific (NSS) *Mutra* splice variants are indicated with arrows. ‘Ov’ refers to ovary samples and ‘Bo’ refers to body samples without ovaries. 100bp ladder (Thermo Fisher) indicates fragment size. Fragments were visualized on 1.5% TAE agarose gel stained with Midori Green (NIPPON Genetics).

