## Supplementary figures and images for "Evidence for an one-step mechanism of endosymbiont-induced thelytoky in the parasitoid wasp, *Muscidifurax uniraptor*"

### Supplementary Figure 1

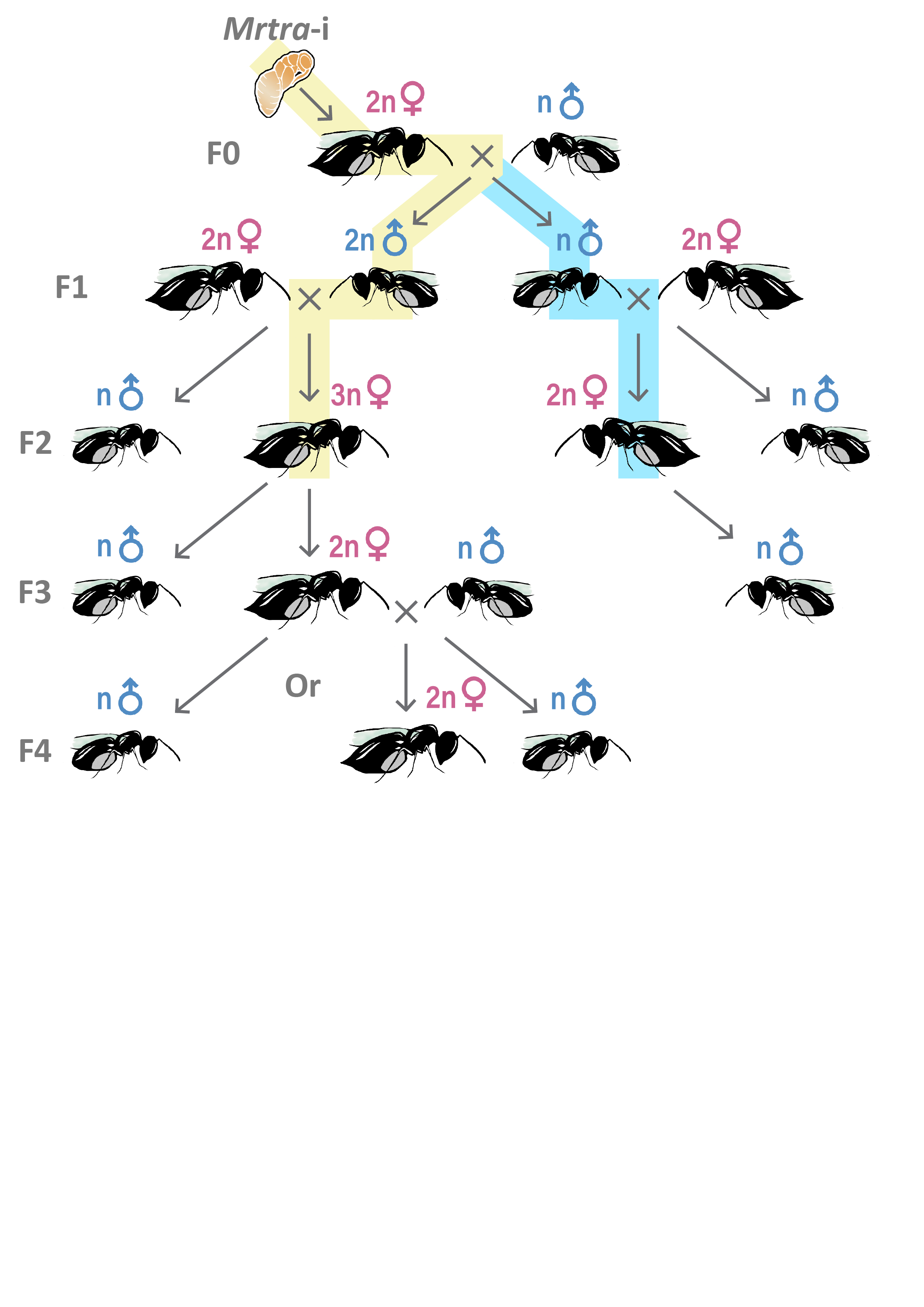

### Supplementary Figure 2

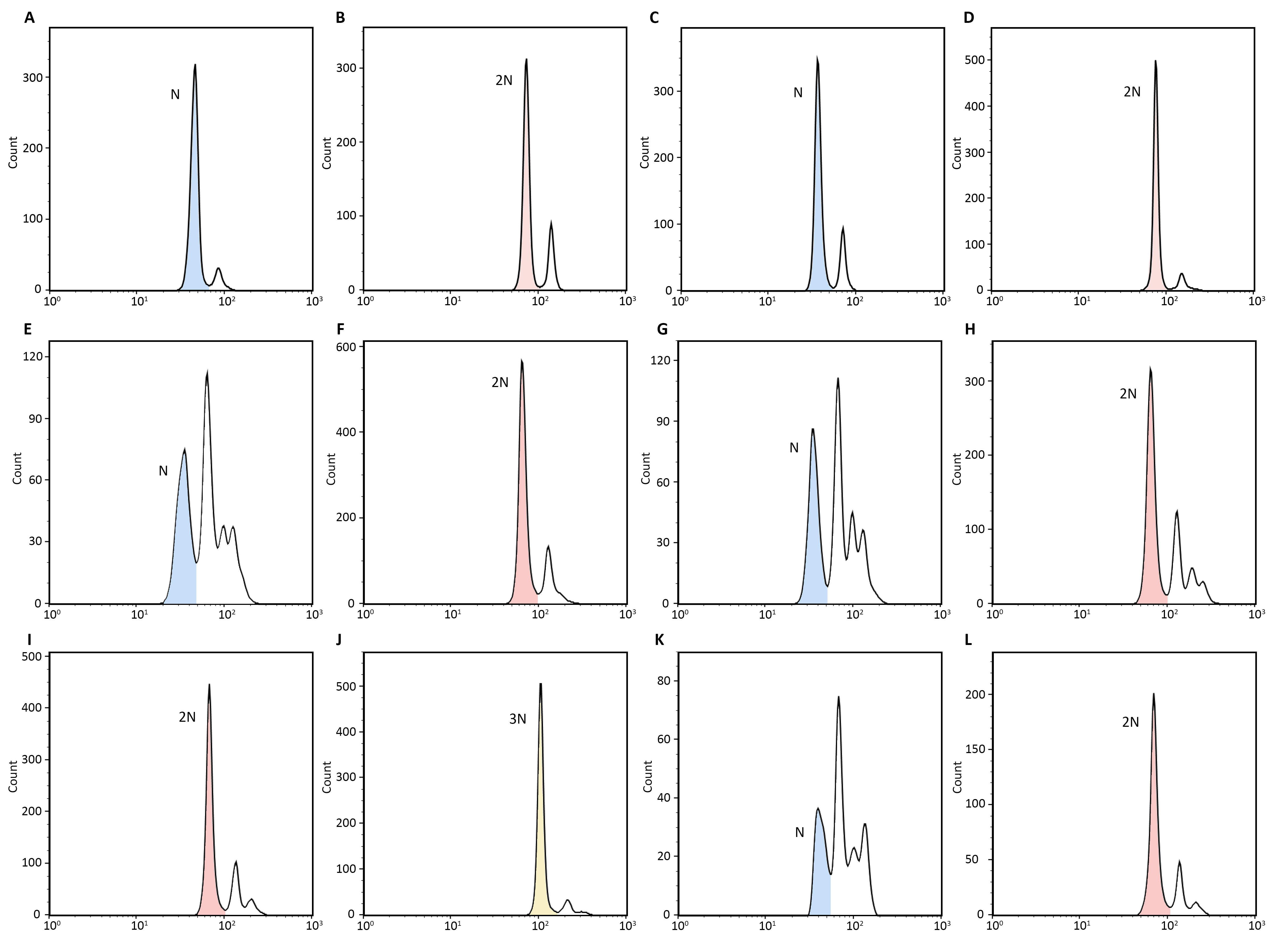

### Supplementary Figure 3

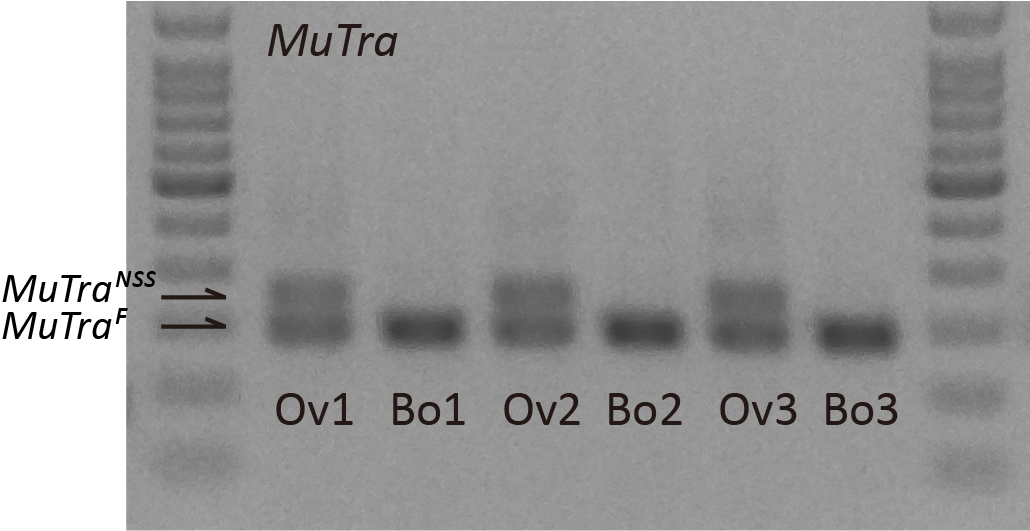
